# Nonhuman primates exploit the prior assumption that the visual world is vertical

**DOI:** 10.1101/2022.10.28.514095

**Authors:** Mohammad Farhan Khazali, Nabil Daddaoua, Peter Thier

## Abstract

When human subjects tilt their heads in dark surroundings, the noisiness of vestibular information impedes precise reports on objects’ orientation with respect to earth’s vertical axis. This difficulty is mitigated if a vertical visual background is available. Tilted visual backgrounds induce feelings of head tilt in subjects who are in fact upright. This is often explained as a result of the brain resorting to the prior assumption that natural visual backgrounds are vertical. Here, we tested whether monkeys show comparable perceptual mechanisms. To this end we trained two monkeys to align a visual arrow to a vertical reference line that had variable luminance across trials, while including a large, clearly visible background square whose orientation changed from trial to trial. On around 20% of all trials, the vertical reference line was left out to measure the subjective visual vertical (*SVV*). When the frame was upright, the monkeys’ *SVV* was aligned with the gravitational vertical. In accordance with the perceptual reports of humans, however, when the frame was tilted, it induced an illusion of head tilt as indicated by a bias in *SVV* towards the frame orientation. Thus all primates exploit the prior assumption that the visual world is vertical.

## Introduction

Our perception arises from the integration of sensory information about the environment with prior knowledge. Underpinning the processing of sensory information with expedient prior knowledge may substantially improve the quality of perceptual decisions (Clemens et al., 2011; Ma, 2019; Vingerhoets et al., 2009). Spatial judgements, dependent on vestibular and proprioceptive signals, are a case in point. Relying on these signals is not only compromised by inevitable sensory noise but also by an inherent ambiguity due to the equivalence of gravitational and translational linear acceleration (Laurens et al., 2013). As a consequence of these limitations, judgments on head orientation in darkness, in which only vestibular and proprioceptive information are available, are unprecise(AUBERT, 1861; MÜLLER, 1916). This is different if vision of the environment provides a reliable frame of reference from which judgements are anchored. The orientation of the observer relative to the outside world can be judged using knowledge that key structural elements of the retinal image of the visual world like the horizon, trees, buildings and the like are well-aligned with the vector of gravity (**Fig 1A and B**). In other words, vision opens access to prior information. However, in order to use this access, the visual system has to figure out the orientation of the image of the visual world with respect to the head. This requires the measurement of its rotation relative to the cardinal axis of the retina and the consideration of the amount of ocular counterroll of the eyes relative to the head. Information on ocular counterroll and other torsional eye movements –typically small in humans– can be drawn from a combination of proprioceptive and efference copy information (Daddaoua et al., 2008; Khazali et al., 2020) whereas the amount of image rotation is gauged by the visual system. Arguably, the precision of this visual estimate might benefit from the overrepresentation of vertical and horizontal structures in the image that should allow noise reduction by pooling (Betsch et al., 2004; Girshick et al., 2011). The dominance of the prior on world orientation provided by the visual background is clearly documented by the counterfactual perception of feeling tilted when exposed to tilted backgrounds, overriding the vestibular report of being upright (**Fig 1C**) (Li & Matin, 2005; Witkin & Asch, 1948).

**Figure 1.**
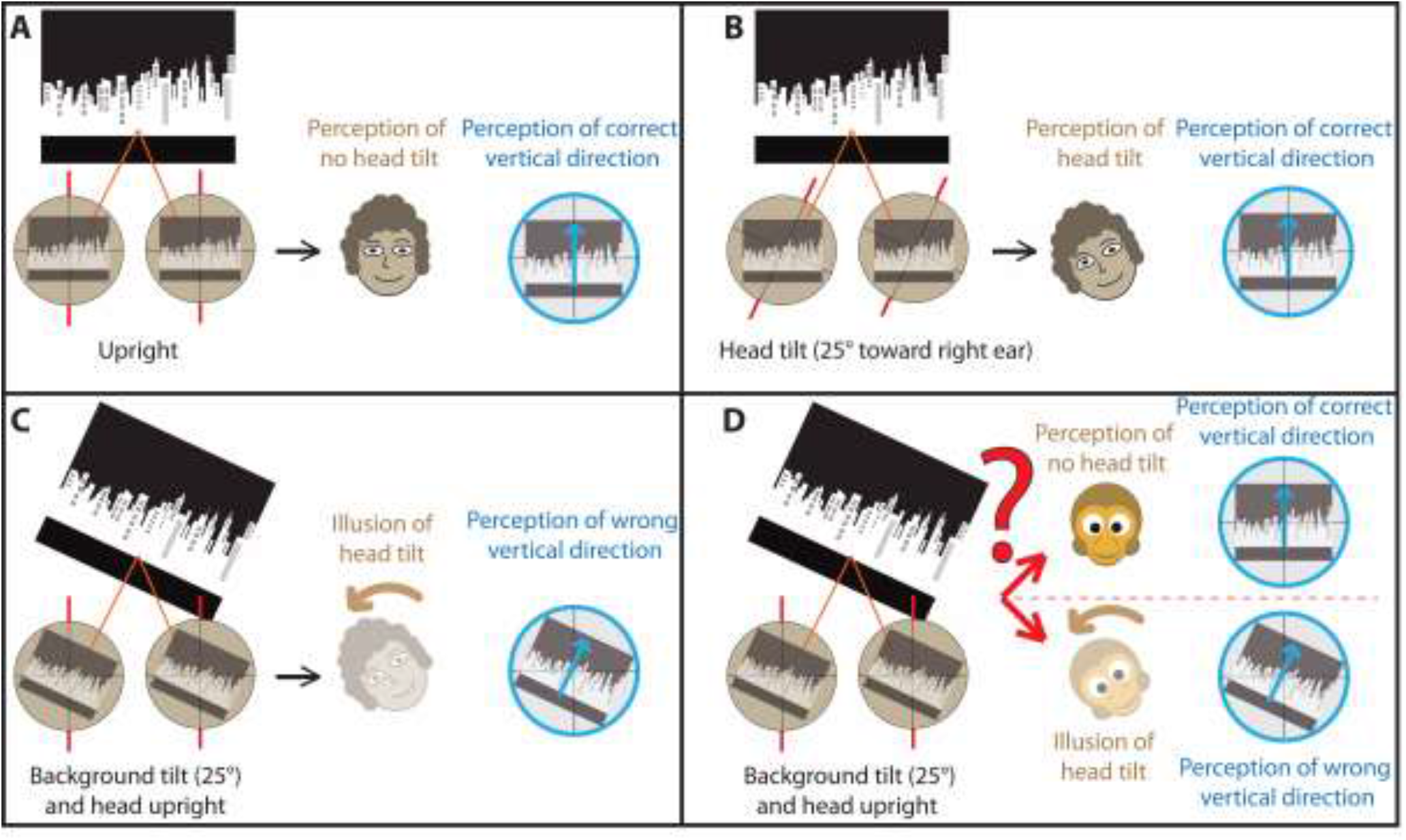
illustration of the hypothesis. (A) shows daily experience when humans experience upright head position, aligned with the orientation of the visual world. Note the brown filed circles represent the retina with its vertical and horizontal axes. The red axes under the retina represent the head main axis, and we ignored here the eye counter roll movement for simplicity. The world image on the retia is representing the orientation of the world with respect to the retina. (B) when the head is tilted 25°, the orientation of the world image on the retina rotates to the opposite direction, but the perception stays stable with respect to the world image orientation. (C) shows basis of the illusion of tilt in humans as the world appears tilted with respect to the head. (D) same as C but in monkeys.

Here, we examine whether rhesus monkeys also use prior knowledge on the orientation of the visual world to judge their orientation (**Fig 1D**). The common ancestors of humans and monkeys lived in a visual world structured by gravity making it likely. The required access to the subjective state of a non-verbal primate species like the rhesus monkey (Macaca mulatta), however, has made verifying this perceptual continuity in the primate line difficult.

In order to finally clarify if the tilt perception of monkeys depends on judgements on orientation of the visual world, we trained two rhesus monkeys on a task measuring the subjective visual vertical (*SVV*) in the presence of a visual background. In this task the animals had to report the orientation of a missing line, embedded in a series of oriented lines of varying visibility. Decisions on the orientation of the missing line served as a measure of *SVV*. The *SVV* of both monkeys coincided with the true vertical when tested against a large visual frame as proxy of a structural visual world, whose vertical axis was upright. However, when the frame got tilted, the monkeys exhibited reports in accordance with the human head tilt illusion, indicated by a bias of their reported *SVV* towards the frame tilt. We therefore conclude that also non-human primates resort to prior assumptions on the orientation of the visual world that can be modelled as the Bayesian combination of a visual prior and a vestibular afference.

## Methods

### Subjects

We used two rhesus monkeys (*Macaca mulatta* M1, M2) in our experiments. Head movements were prevented by deploying a chronically implanted head post that enabled easy and painless stabilization of the head. Head post-surgery was carried out under intubation anesthesia with isoflurane (0.8%), supplemented by continuous infusions of Remifentanil (1–2.5 μg/kg min) and tight monitoring of all relevant parameters (body temperature, heart rate, blood pressure, pCO_2_, and pO_2_) that allowed prompt and adequate reaction in case of deviations from normal. Buprenorphine (0.01 mg/kg body weight twice a day) was given to eliminate post-operative pain. Analgesia was stopped as soon as the animal had returned to its normal pre-operative pattern of behavior, a change we took as indication of the disappearance of pain. All surgical procedures complied with the NIH Guide for Care and Use of Laboratory Animals and were approved by the local animal care committee.

During an experimental session, the monkeys were seated in a primate chair while the aforementioned head post was fixed with respect to the chair using a head holder attached to the chair. They turned a lever of 10 cm length located at the back of the front panel of the primate chair left or right to move an arrow to align it with the direction of verticality within the framework of the behavioral paradigm described below.

### Behavioral setup and training

The visual stimuli were presented on a fronto-parallel LCD screen (19 in., resolution: 1280 × 1024) located at 38 cm in front of the monkey. The monkey watched the screen binocularly through a circular aperture (40.5° visual angle) preventing him from using any of the screen edges for orientation. The primate chair and the screen were surrounded by an opaque sphere (diameter 196 cm) dimming out the room around the monkey and effectively eliminating any visual cues that might have served as orientation landmarks.

In the experimental sessions (**Fig 2**), the monkeys saw a visual line of 40° length and 2° width centered in the aperture. It served as reference indicating the gravitational vertical. Whenever this line was presented together with a squared visual frame (28°x28°, line width of 3°), its length was chosen such as not to exceed the frame borders whatever the tilt angle was. Depending on the orientation of the frame the reference line length varied between 28° and 40°. In *training trials*, which had a share of 80%, the contrast of the reference line was randomly chosen from a fixed set of values (see below). In complementary test trials, randomly interspersed, the reference line contrast was set to 0%, hence rendering it invisible. The reference line contrast level was given by the Michelson definition: contrast level (in %) = 100 × (reference line luminance − background luminance) / (reference line luminance + background luminance). In addition to the reference line, a white arrow (length: 26.5°; width: 1.5° visual angle; contrast 95%) was presented, whose orientation in the roll plane could be adjusted by the monkey with an angular resolution of 0.6°. The edges of the reference line, frame, and the arrow were anti-aliased by the experiment software written Python2.4 and the Vision Egg interface to keep the arrow shape constant during rotation(Freeman, 1974; Straw & Perrinet, 2008), thereby ensuring that the monkey would not use individual pixels changes as basis for judgments on the orientation of the arrow. The monkey controlled the orientation of the arrow by turning the lever. Turning the lever by 20° to the left closed an electronic contact, associated with a mechanical stopper, which led to a continuous counterclockwise rotation of the arrow at a constant velocity of 10 °/s. Correspondingly, when the monkey turned the lever by 20° to the right, the arrow started to rotate continuously in a clockwise manner at the same constant velocity. As soon as the monkey moved the lever to a position in between the two extremes, the rotation of the arrow stopped. During the *training trials*, the monkey received a reward only if the angular deviation of the arrow from the reference line fell below a pre-set threshold. Initially, the acceptable deviation was big (±25°), but in the course of the training, gradually reduced to ±3°. Note that the monkey had only one chance per trial to meet the threshold criterion as no corrective adjustments of the arrow were allowed. If the arrow stopped outside the acceptable range, the monkey had to wait for the next trial in order to get a new chance. During the experiment (**see Fig 2**) the monkeys were asked to align the arrow to the direction of gravity independent of the presence of the squared frame (length: 28° visual angle of each side) at 95% contrast level. The orientation of this frame varied across trials and in some it was absent.

**Figure 2.**
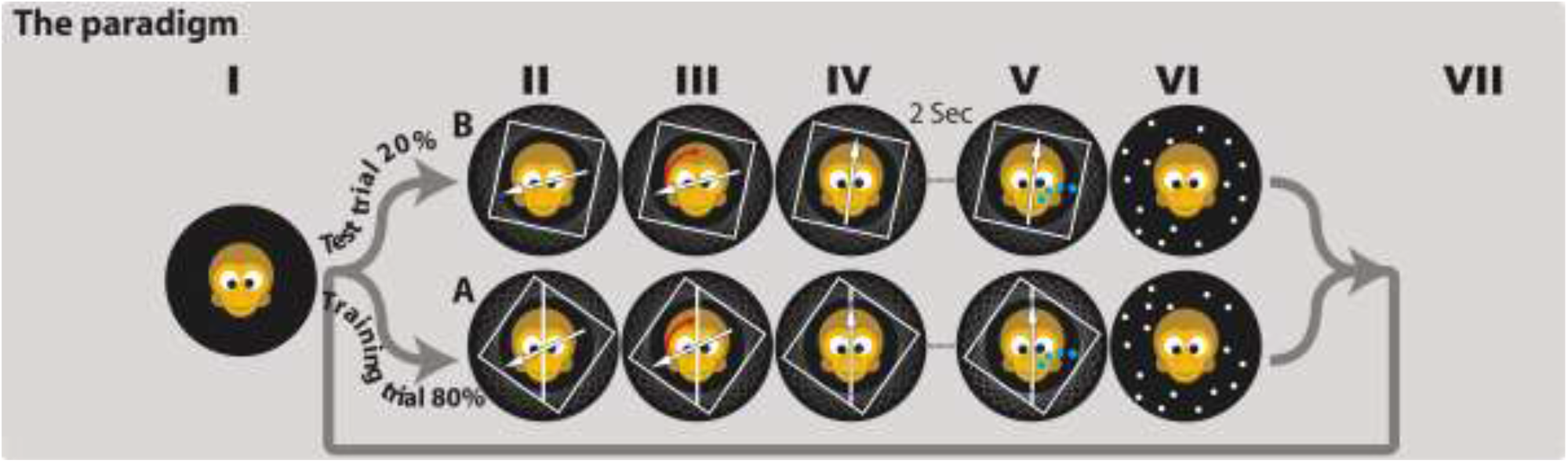
the paradigm. (I) start the experiment with darkness. (II) Probing the *SVV* started 30 seconds after. The frame was presented at 0°, ±12.5°, ±22.5°, ±35°, and ±45° tilt angle with respect to the gravity vector. (III-IV) The monkeys were asked to move the arrow to the desired position. (V) They were rewarded if they aligned the arrow correctly (*training trials*), and randomly rewarded for *test trials* independent on the arrow alignment. (VI) A random dotes pattern was presented for 1 second to erase potential after effects of the visual stimuli used. (VI) The end of the trail and initiating a new one.

### Experimental paradigms

During the experimental sessions, the monkey was exposed to a series of trials chosen at random from two classes: *training trials* and *test trials*. In the *training trials* (which constituted 80% of all trials for M1 and M2), the monkey was asked to orient the indicative arrow upward by aligning it as precisely as possible with the visible vertical reference line whose contrast could be chosen from a fix set of values (93.5 %, 89 %, 79 % and 50 %, see **Fig 2A**). The initial orientation of the arrow was randomized. The orientation of the arrow adopted, when its movement stopped for the first time, was considered as the monkey’s choice. The monkey was rewarded with a drop of apple juice if the angular deviation between the arrow and the reference line fell below 3°. In the *test trials* (which constituted 20% of all trials for M1 and M2), the contrast level of the reference line with respect to the black background was set to 0 %–i.e. it was effectively absent (see **Fig 2B**). A reward was provided at random in 70% (M1) and 50% (M2) respectively of the *test trials* if the monkey stopped turning the arrow at any position after the beginning of the trial, irrespective of the angle chosen. Irrespective of the trial type a squared frame was clearly visible (95 % contrast) and its orientation, with respect to an upright square, was chosen randomly from a fix set of values (0°, ±12.5°, ±22.5°, ±35°, ±45°, the minus sign designating counterclockwise roll rotations and the plus sign clockwise roll rotations from the monkey’s point of view) from one trial to the next.

### Modeling

In an attempt to use the behavioral decisions of monkeys to estimate their perception of head tilt, we resorted to a simplified version of a Bayesian model proposed by Vingerhoets et al. (2) to model human perception. It predicts the subjective head orientation based on a combination of noisy vestibular signals on the amount of true head tilt and the prior assumption that the head is aligned with the vector of gravity. This assumption decreases the amount of variability of head tilt estimates at the expense of introducing a bias towards upright. The original model considers another assumption about the background of the visual world, which is that it is always vertical. It is able to estimate the perceived amount of head tilt for different objective head tilts, carried out in darkness or in the presence of a visual squared frame (De Vrijer et al., 2008; Vingerhoets et al., 2009). In our case, there was no objective head tilt but an illusion of head tilt induced by the tilted frame (i.e. the visual background). Hence we skipped the original model’s premise that the head is aligned with the vector of gravity while keeping the assumption that the orientation of the visual background is aligned with the gravity vector.

In this model, vestibular and proprioceptive inputs are captured by the likelihood function 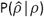 that specifies the estimate of head tilt 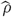 given the objective head tilt *ρ*. The quantitative characteristics of the likelihood function were modelled as a normal Gaussian distribution centered at the objective head tilt *ρ* and the standard deviation *std*_*ρ*. In the absence of objective head tilt, it should ideally be centered on zero, i.e. parallel to the vector of gravity. Assuming that the *SVV* as measured in darkness should only depend on the vestibular and proprioceptive inputs, the standard deviation *std*_*svv*of the *SVV* can be taken as estimate of the model distribution, *std*_*ρ*, which reflects the noise in head tilt inputs. A slight shift of the *SVV* away from zero is a manifestation of a bias. For M1 the *SVV* was -3.0° with *std*_*svv* equaling 9.5°, whereas in M2, the *SVV* was - 0.59° with *std*_*svv* equaling 5.8°. This likelihood function was merged with a visual likelihood function representing the contribution of the tilt of the retinal background image to the perceived head tilt. Vingerhoets et al. (Vingerhoets et al., 2009) described the visual likelihood orientation function as a Von Mises distribution– a distribution close to a Gaussian distribution for periodic functions–with four equally probable cardinal directions representing the effect of a square frame. The visual likelihood orientation function equals 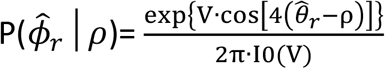 with 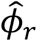, the estimate of retinal image rotation, given the true head tilt *ρ*, under the observer’s assumption that the background is always vertical. 2π serves as a normalizing factor, V is a concentration parameter that equals the inverse of the variance, where I0 denotes the modified Bessel function with the order zero. To keep the model manageable we refrained from considering expansions to account for the possibility of better resolution for horizontal and vertical orientations as suggested by (Girshick et al., 2011).

The model’s main assumption is that the orientation of the background (i.e. the squared frame) is aligned with the gravity vector. Therefore in the absence of non-visual information on head tilt, the estimated vertical direction become ambiguous because it is the same if the frame is upright or rotated by 90°, 180° and so on (Li & Matin, 2005). Whatever the frame effect might be, it should repeat itself four times in 360° of frame rotation (EVANS M, H. N., 2000; Girshick et al., 2011; Vingerhoets et al., 2009). What resolves this ambiguity is the merger of information provided by the frame orientation and vestibular and proprioceptive information on the head tilt. The visual ambiguity is a consequence of the fact that the frame could rotate either in a cw or a ccw direction, given the rotational symmetry of the frame with the frame tilted 45° cw or 45° ccw (and equally 45° plus multiples of 90°) being perceived as identical. Hence, in the absence of a decision bias, the observer will provide equal numbers of cw and ccw decisions, the two cancelling each other out suggesting a *SVV* that does not deviate from the true vertical. On the other hand, for frame tilts between 0 – 45° cw, the *SVV* will be perceived as rotated in a cw direction and conversely rotated in a ccw direction for frame tilts between 0 – 45° ccw. Finally, because of the rotational symmetry we expect that the frame effect to repeat itself four times in 360° of frame tilts, a pattern that was clearly exhibited by our data and captured by the visual likelihood orientation function.

We used Bayes’ rule to calculate the posterior probability distribution 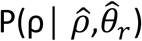 results from merging the frame orientation likelihood function 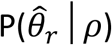 and head tilt likelihood function 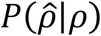 resulting in 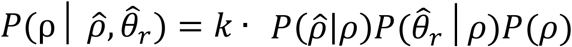, hereby keeping in mind that *P*(*ρ*) =1as we did not have any prior knowledge about the head orientation for simplicity, allowing us to decrease the model’s free parameters. *K* is a normalizing constant for the posterior probability. The peak of this distribution is the maximum posterior probability (MAP) that indicates the subjective head orientation in space (*β*). Based on the value of *β*, the monkey will align the arrow by the same amount but in the opposite direction, deviating the arrow orientation on the retina 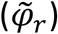. If *β* is zero then 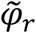 will be zero too and the linearly summation of both will represent the line orientation in space (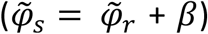, which is ideally zero, in this case manifesting the perception of no head tilt. However, an illusion of head tilt due to frame rotation, will result in a deviation of *β* from zero (e.g. 5° cw). Since the monkey tries to align the arrow as vertical as possible based on his perception 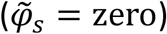, the arrow will deviate from the retinal meridian by 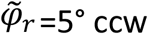 ccw. Thus when the monkey reports 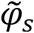 deviation from zero, this indicates the perception of head tilt. The model fitting explains the *SVV* modulation as function of the squared frame tilt very well, see (**Fig 5**), having just two free parameters, the vestibular likelihood function center and V, the concentration parameter from the frame orientation likelihood function.

### Fitting procedure

We fitted our data to the model described above to get the parameters for the best fit using the least square method provided by the Matlab R2011b routine fminsearch. We used the measured *SVV* in darkness (no frame) to obtain the center *ρ* and standard deviation *std*_*ρ* of the head tilt likelihood function 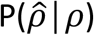. During fitting we kept *std*_*ρ* constant since it represents the vestibular and proprioceptive information independent of the influence of the frame. The center of the likelihood function (*SVV* in darkness) was used as an initial value for the frame effect offset. V was the only free parameter representing the frame orientation likelihood function. The fitting was excellent for both monkeys as shown in **Fig 5**, yielding very small sum of square errors of 5.1° and 11.3° and r^2^ values of 0.89 and 0.51 for M1 and M2 respectively.

## Results

We trained two monkeys (M1 and M2) to report their *SVV* by aligning an arrow, starting from a random orientation, to a vertical reference line whose visual contrast was varied across *training trials* (see **Fig 2A** and **Methods)**. The aim of the *training trials* was to develop an association between the orientation of an indicative arrow, to be adjusted by the monkey, to the orientation of the vertical reference line serving as proxy of the concept of vertical (the subjective visual vertical, *SVV*), available even in the absence of the reference line. The existence of this association was measured with interspersed *test trials* in which the reference line had a contrast level of 0%, i.e. was rendered invisible(Daddaoua et al., 2008). *Test trial* had an overall share of 20% (see **Fig 2B**). In order to test the impact of a visual background on the *SVV* we resorted to a squared frame as a proxy for the visual background, a simple, abstract pattern known to induce a tilt illusion in humans (Ebenholtz & Glaser, 1982; Li & Matin, 2005; Wenderoth & Beh, 1977). Arguably such patterns should not have meaning for a monkey. Hence, they do not come with the danger that a monkey’s response strategy might be biased by a semantic interpretation of the background. Rather, the perceptual impact of this abstract pattern may be safely led back to the presence of the vertical and horizontal orientations that also dominate natural visual backgrounds.

Once the monkeys had understood the need to align the indicative arrow with the reference line, we could use its *test trial* orientation as a manifestation of the *SVV*. In order to study the influence of the visual background on the *SVV*, we varied the orientation of the squared frame in a randomized manner from one trial to the next and calculated the mean *SVV* for the various squared frame orientation classes (see **Methods**). The right part of (**Fig 3)** depicts plots of the mean *SVV* as function of the frame orientation in space for the two monkeys in *test trials*. For comparison, the left side shows plots of mean arrow orientation as function of the frame orientation for the same monkeys during *training trials*. Not unexpectedly, the ability to align the indicative arrow with the reference line is practically perfect (mean ± std= 0.17°ccw ± 2.1 for all *training trials* pooled together). We subjected the *training trials* to a two-ways ANOVA with the *reference line contrast level* and *frame tilt* angle as two independent factors. We found no effect of *reference line contrast level* (p =0.44; four contrast levels with n=1215 for each). In contrast, the *frame tilt* affected the *misalignment* significantly (p= 0.008; nine *frame tilts* with n= 540 for each angle). There was no significant interaction between both factors (p= 0.6564; n= 135 for each condition). Thus, even when the arrow alignment to the reference line looked almost perfect by eye, a small amount of systematic *misalignment* was induced due to the *frame tilt*. This *misalignment* peaked at 12.5° cw and ccw with an amplitude of 0.17 and 0.28° in cw and ccw directions respectively. These *misalignments* are measured with respect to the *misalignment* when the frame was vertical (mean= 0.16° ccw). Thus, the effect of the tilted frame on the monkeys’ perception seems to be natural and strong enough to influence the alignment of an arrow to a clearly visible reference line. This unexpectedly clear influence of the frame suggested a much stronger impact of frame orientation on the *SVV* in the absence of the reference line. Indeed the *test trials* showed not only more variable *SVV* but also a clear systematic dependency on the *frame tilt* characterized by a maximal deviation (∽4°) from the true vertical at (12.5-35° cw and ccw) *frame tilt* in the direction of the tilt. The *SVV* was aligned with the true vertical when the frame was vertical or tilted by ±45°. Thus, the frame acts as an attractor for the *SVV*, arguably because it induces an illusion of head tilt in monkeys.

**Figure 3.**
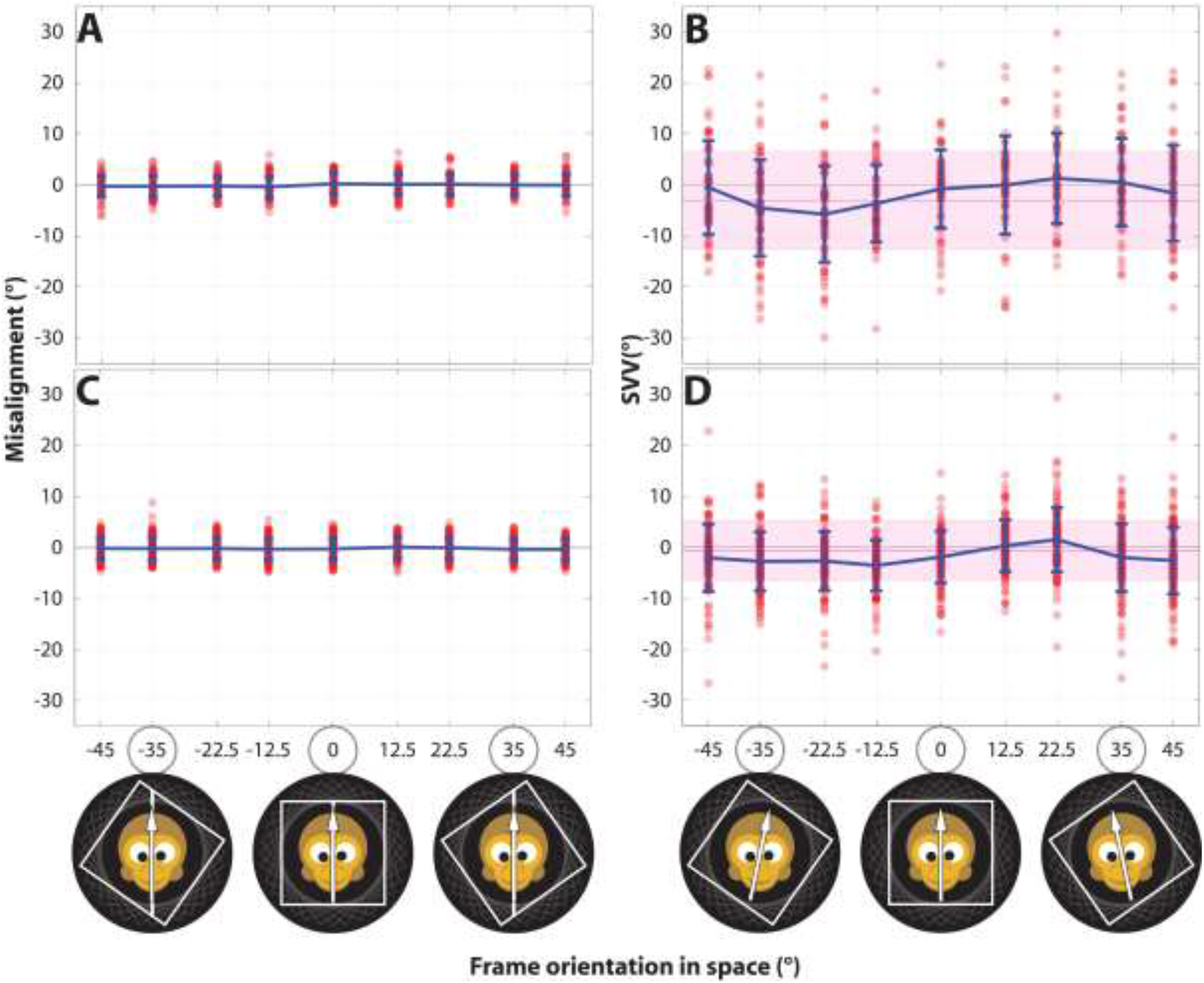
SVV during presenting the background. Plots of arrow *misalignment* to the reference line, in blue, as function of frame-tilt angle for *training trials* (A and C) and it plots the *SVV* in *test trials* (B and D) for the monkeys M1 and M2 respectively from up to down. The vertical bars indicate the standard deviations, and the pink horizontal line indicates the *SVV* in compete darkness, where the shadow indicates its standard deviation and each dote represents the *misalignment* or *SVV* in each trial. Note that M1 and M2 performed 3199 and 5079 trials, respectively.

In order to test the validity of the impression of a systematic influence of the squared frame orientation on the *SVV*, characterized by a deviation from the true vertical, we subjected the *test trials* data to a one-way ANOVA with the factor *frame tilt* angle in space separately for the two monkeys. This analysis indicated a significant effect of the *frame tilt* in space on the *SVV* of both M1 (p=1.4 *e*^−5^) and M2 (p < 1 *e*^−16^). Note that zero SVV for frame tilts of 0° and for +/-45° indicate that the *SVV* would change as a function of the frame orientation, repeating itself every four times in 360°, similar to what has been found in humans (Li & Matin, 2005).

As said, both monkeys reported a *SVV* aligned with the true vertical when the frame was upright (i.e. its edges having horizontal and vertical orientations respectively) or when tilted 45°. When the frame was upright, the *SVV* accuracy – i.e. the match between the *SVV* and the true vertical - was significantly better compared to the *SVV* accuracy in the absence of a frame in monkey M1 (no frame: mean ± std = -3.0±9.5°; with frame 0.45±4.1°; t-test, p=0.006). Monkey M2 had already a very accurate *SVV* in the absence of a frame, which did not provide a large margin for further improvement (no frame: -0.59±5.8°; with frame-0.66±4.2°; t-test, p=0.91, **Fig 4 left panel**).

**Figure 4.**
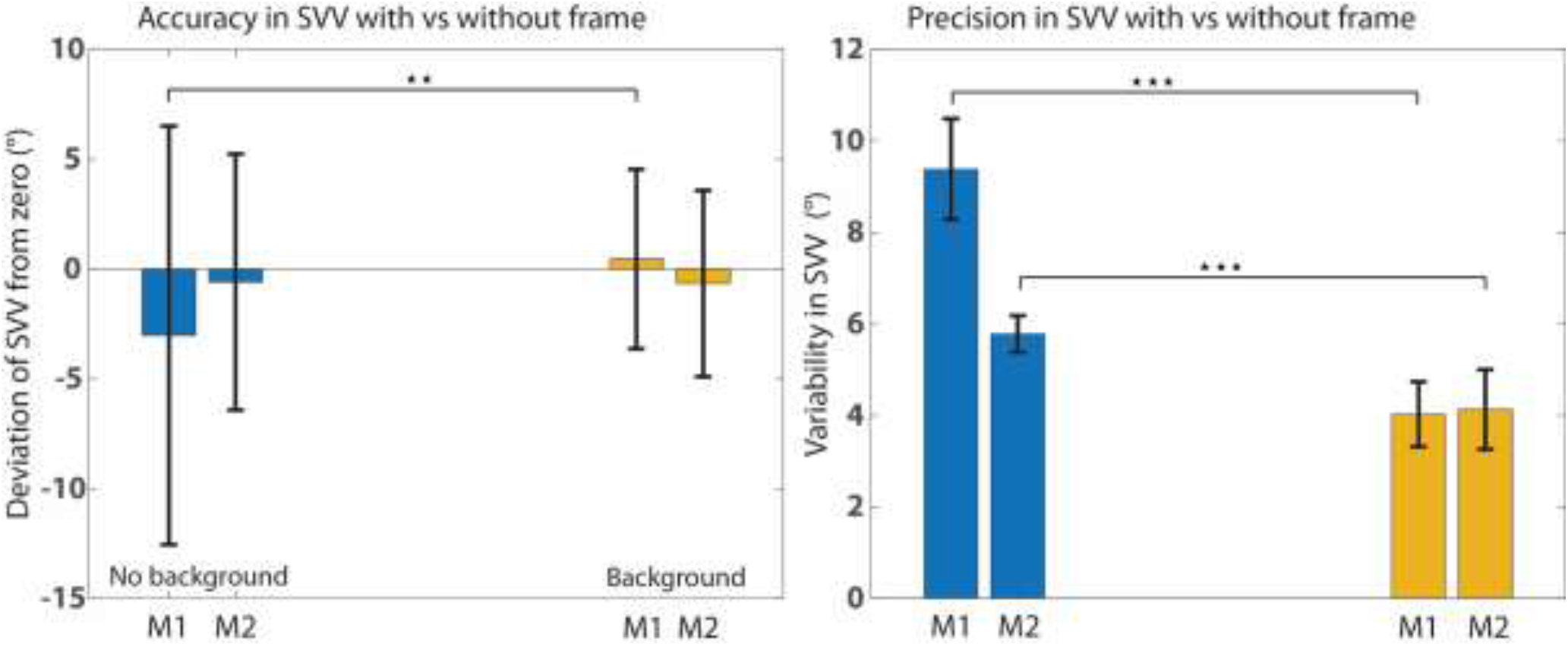
enhancing accuracy and precision by presenting vertical frame. Left panel is a plot of mean deviation of *SVV* and its standard deviation, in blue during darkness (no frame) and in yellow during presenting vertical frame. The right panel shows the precision of the measured *SVV* in blue for during darkness an in yellow during upright frame. Note that both the accuracy and precision improve when an upright frame is presented. The right panel was plotted based on bootstrapping technique. Basically we calculated the standard deviation 1000 times from randomly chosen sample points with return from the data of the same size of the original data. Variability in the *SVV* plotted on the y axes is the mean of these 1000 standard deviations.

We next examined whether the frame improved the precision of the *SVV*, i.e. the variation of *SVV* reports, by resorting to a bootstrapping approach. We randomly drew with replacement the same amount of trials, collected for each condition and each monkey (altogether n=71 and 119 with frame; n= 61 and 104 without frame, for M1 and M2 respectively). The trials were collected over 41 experimental sessions with 16 sessions for M1 and 25 sessions for M2. The bootstrapping procedure was repeated 1000 times, allowing us to derive 1000 individual *SVV* estimates as basis of calculating a mean *SVV* and its standard deviation, the latter as the measure of the *SVV* precision. This then allowed us to compare the precision of *SVV* reports across the two conditions. In both monkeys the standard deviation (std) decreased – i.e. the precision increased –from the no frame to the frame condition (mean (std) ± std (std) = 9.4±1.1° for M1; and 5.8± 0.4° for M2) to (4.0± 0.71° for M1 and 4.1± 0.87° for M2), respectively (p <0.001; t-test two samples; for the pooled data from both; **Fig 4 right panel**).

Finally, we tried to interpret the influence of the frame as a consequence of an optimal integration of different streams of information in a simple Bayesian framework. Similar to other Bayesian models trying to capture key principles of perception, also our model assumed that various sensory modalities are integrated with prior knowledge (De Vrijer et al., 2008; Vingerhoets et al., 2009). For the sake of simplicity, we restricted our model to situations in which the head was actually upright, i.e. aligned with the gravitational vector in accordance with the conditions explored in our experiments. We did not consider any premises as to the orientation of the head. Note that the subjective experience of upright was given by *SVV* measurements with no frame (see **–Modeling in Methods** for more details) that could reveal slight deviations from the true physical vertical, reflecting an internal bias that had to be taken into account when centering the likelihood function of the vestibular and proprioceptive signals. Merging the frame orientation and the head tilt likelihood functions allowed the model to fit the *SVV* data as a function of the frame tilt very well, yielding small sum of square errors of 5.1° and 11.3° and r^2^ of 0.89 and 0.51 for M1 and M2 respectively (**Fig 5**). The fitting was done with two free parameters, the first the frame effect offset = -1.5° and -1.0° and the second the concentration parameter of the Von Mises function (V) = 0.60 and 0.33 for M1 and M2 respectively. The *SVV* fit exhibited the expected periodic profile that repeated itself four times in 360°. As discussed earlier, the periodicity is a result of the interaction between a symmetric square frame that repeats its orientation four times in 360° and the ambiguity regarding the direction of rotation of the square. The *frame tilt* impact peaked at (12.5-35° cw and ccw) relative to upright and was zero for tilts of (45° cw and ccw). Thus our Bayesian approach to model the integration of these multiple sources of information indicates why even the consideration of a simple background like the squared frame is still advantageous, when it is truly vertical reflecting a proxy condition of the real world.

**Figure 5.**
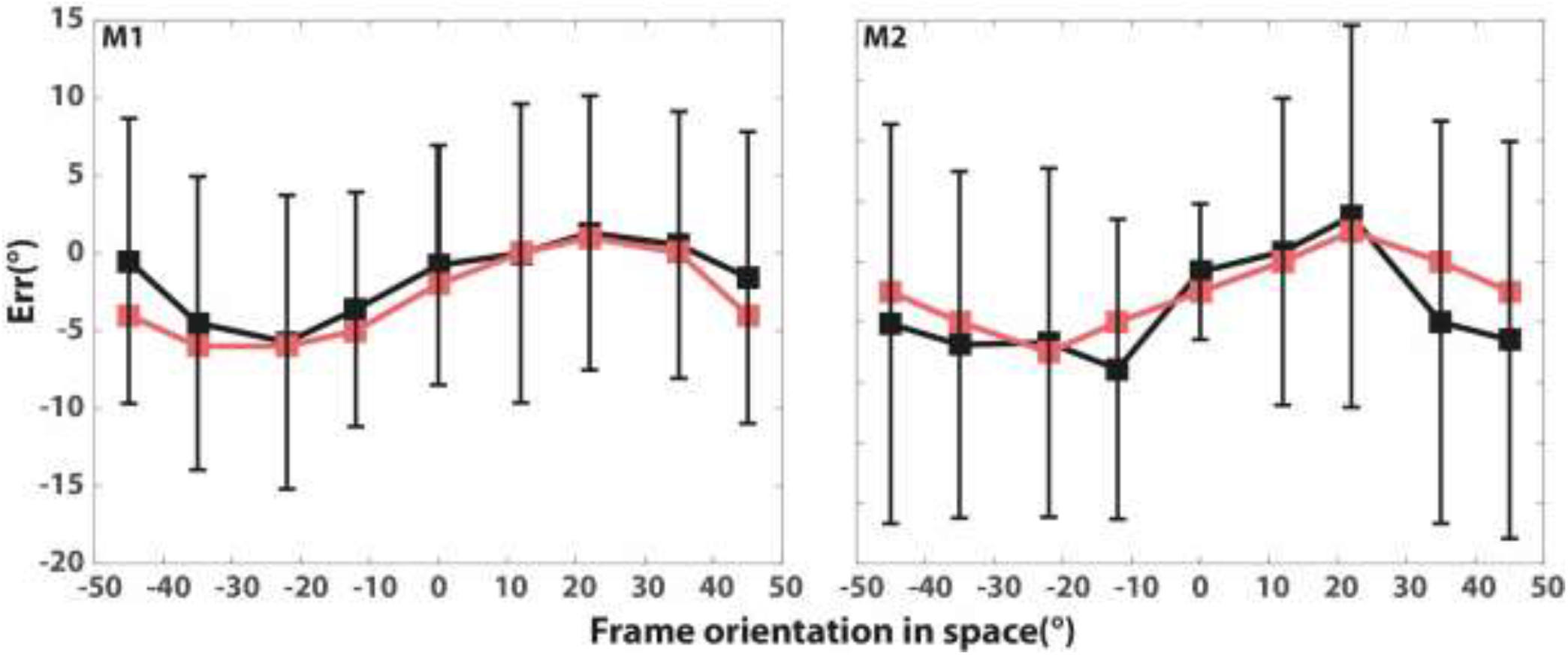
modeling the SVV as a function of the frame tilt. *SVV* as a function of the frame orientation in space with their standard deviation in black, and the Bayesian model fitting is in red. The left panel is for monkey M1 and the right one is for M2. The fitting was done having two free parameters, the first was the frame effect offset = -1.5 and -1.0 for M1 and M2 respectively and the second was the concentration parameter (V) = 0.60 and 0.33 for M1 and M2 respectively, see **Methods** for more details.

## Discussion

In this study we were able to show that rhesus monkeys exhibit an illusion of head tilt, despite being upright without physical head tilt, when they are confronted with a tilted visual background. The assumption of a tilt illusion was grounded in behavioral decisions that matched the expected influence of the illusion. Our monkey observers had been trained to align an indicative arrow with the vertical axis by associating the arrow with a clearly visible vertical reference line. Assuming that the concept of the vertical axis thus rehearsed would be sustained in occasional *test trials* lacking the reference line. We used the influence of a tilted squared background frame on the orientation of the indicative arrow as measure of its influence on the *SVV*, applying the same approach as Daddaoua et al.(Daddaoua et al., 2008). This allowed us to use the influence of a tilted squared background frame on the orientation of the indicative arrow as measure of its influence on the *SVV*. We are confident that this is indeed a valid assumption for the following reasons: First, the monkeys adjusted the indicative arrow close to the gravitational vertical in darkness, when neither the reference line nor a background were available. Second, the deviation of the *SVV* prompted by the frame was systematic and consistent for both monkeys. Third, the frame tilt effect was very small, not exceeding a peak deviation of ∽ 4° for frame tilts around 12.5-35°, either cw or ccw. Assuming that such subtle, yet consistent and systematic effects might be intentions reflecting unknown idiosyncratic preferences of the experimental subjects is neither parsimonious nor epistemologically promising. Fourth, the frame tilt effects documented by us are very similar to the ones found in humans, which until now were unable to detect because monkeys are not able to verbalize their subjective experiences. And fifth and finally, the approach we used to train the monkeys to indicate the vertical direction had already been successful in measuring the *SVV* when roll tilting the head tilt in monkeys (Daddaoua et al., 2008). Hence, we are confident that our approach to gauge our monkeys’ perception of the vertical and its modulation by a visual background led to meaningful results that document a clear continuity in the primate line.

The monkeys’ *SVV* was close to the true vertical without additional visual cues, similar to the monkey’s study, mentioned above, which measured *SVV* in darkness while roll tilting the monkey’ s body and head (Daddaoua et al., 2008). The results of the overlapping conditions between both studies, namely reporting the *SVV* in an upright position and in darkness were similar, close to the true vertical, providing additional evidence that the approach chosen in this and previous work indeed captures the *SVV* of monkeys. Finally, our results show very similar effects of the background on the monkey and on the human SVV (Vingerhoets et al., 2009). Presenting a vertical frame benefited monkey M1’s perception of the vertical by reducing the deviation of his *SVV* from the physical vertical by almost two degrees. Monkey M2 had already a very small *SVV* deviation of only ∽0.5°, which did not offer much room for further improvement. Both monkeys also showed a higher precision of their *SVV* when a vertical frame was presented. Finally the *SVV* exhibited a cyclic deviation from the true vertical toward the frame tilt orientation, causing a retinal image orientation that would also be evoked by a true head tilt in a direction opposite to the frame tilt (Li & Matin, 2005; Vingerhoets et al., 2009; Wenderoth & Beh, 1977; Witkin & Asch, 1948) and in the condition discussed here arguably associated with the experience of an illusory head tilt. The frame used in our experiments models natural backgrounds whose retinal images will usually move and adopt new positions only as a consequence of ego motion. An example are head tilts that will cause a counter-rotation of the image of the background. Visual backgrounds are dominated by structures that like trees or buildings are aligned(Betsch et al., 2004; Girshick et al., 2011) with the gravitational vertical and therefore – in the absence of head tilt –also aligned with the vertical retinal meridian. Hence, image tilt relative to the meridians offers valuable information on head tilt that complements information provided by other modalities, thereby allowing more precise and more reliable judgements. That this is indeed the case is documented by the fact that the results obtained in the present study could be predicted by a Bayesian model that accommodates the integration of different sensory modalities and prior information in a statistically optimal manner, able to minimize the impact of noise and thereby improving the precision and reliability of perceptual decision. Note that our model had two roots. The first one was an estimate of the compound contributions of vestibular, proprioceptive and prior information to head tilt perception provided by measurements of the SVV in darkness as proxy of perceived head tilt in the absence of vision. The second one was an estimate of the impact of visual information gained in an experiment that tested a conflict between a visual input indicating head tilt and the other carriers of information denying it. Multisensory integration is in principle beneficial as it will reduce the uncertainty of perceptual decisions. However, it comes with the prize of biasing the decision in the direction of the signal with lower noise. This is very well documented by the results of the present conflict study, in which visual information advocating head tilt was confronted with non-visual information denying it. The results exhibited a significant impact of vision, reproduced in the model under the premise of comparatively low visual noise. This dominance of vision, indicated by a larger peak to peak of *SVV* deviation elicited by tilting the background image was larger for M1 than for M2. This means that visual information had more influence on M1’s decisions, an observation that was captured by a larger concentration parameter V in the Von Mises function and by larger noise in the vestibular and proprioceptive signals for M1 as compared to M2. This is in line with the Bayesian explanation as M1’s *SVV*, which exhibited larger noise ratio from visual to vestibular signals, was associated with larger visual signal influence, in contrast to M2.

Under natural conditions, the impact of the background orientation may in fact have an even stronger effect as to what we found in the present experiments in which we used a large squared frame as proxy of a natural background. In a recent study on humans, were exposed to tilted renditions of their natural visual environment, the bias in *SVV* reached ∽10° for 36° of background tilt (De Winkel et al., 2021). Additionally, our simple geometrical structural not only lacked the rich structure of a natural scene but also came with the inevitable ambiguity of being compatible with in principle four potential directions of gravity due to the frame’s symmetrical shape, an ambiguity that was further increased by the in principle two possible rotation directions. Hence the frame effect was possibly divided by four possible directions.

It may be surprising to see that vision dominates vestibular and somatosensory information in the perception of the vertical. However, this visual dominance is most probably task-dependent, i.e. confined to particular behavioral conditions and requirements. In fact De Winkel et al. (De Winkel et al., 2021) showed that vision dominated human perception of the SVV in tasks in which they were asked to estimate the vertical when exposed to tilted visual backgrounds by aligning a visual rod with the axis of gravity. Yet, when asked to align the platform on which subjects were standing perpendicular to the axis of gravity, thus providing an estimate of the subjective postural vertical, the impact of the tilted visual background was significantly reduced (see (Clemens et al., 2011; Mittelstaedt, 1995) for related findings). The fact that the impact of vision on the subjective visual and the subjective postural vertical differ may be rooted in ecological needs. Obviously, timely vestibular and somatosensory feedback for the control of the biomedical effectors involved are certainly much more reliable than sluggish visual feedback that will become available way too late to prevent imbalance. Moreover, local natural grounds providing support are usually uneven with random deviations from the gravitational horizontal. While these deviations are gauged by the proprioceptive and the vestibular system, only vision is able to assess information on the orientation of wider stretches of the ground not accessible to the other. In fact observations on patients with stroke based lesions suggest that the two combinations of sensory signals on the vertical serving balance as opposed to perception may in fact have different sites in the human brain (Anastasopoulos et al., 1997; Bisdorff et al., 1996; Karnath et al., 2000; Masdeu et al., 2005).

Our results are valuable because they suggest an evolutionary shared ancestry of a key aspect of human and monkey perception. And because of this commonality they may help to bridge the gap between the results of behavioral work in humans and neurophysiological data collected from monkeys. For instance, electrophysiological recordings from the caudal interparietal area of rhesus monkeys have identified neurons that are sensitive to visual background tilts (Rosenberg & Angelaki, 2014), possibly useful as source of information integrated into a yet unknown, putative neural representation of the SVV.

## Conclusion

Non-human primates utilize the statistics of vertical world similar to humans and therefore their perception deviates from true vertical direction when they are introduced to a vertical background. We demonstrated that monkey’s decision about verticality is dominated by visual information provided by the background. Our results fill the gap between electrophysiological knowledge about sensing vertical direction, collected mainly in monkeys, and perceptual studies that were mainly applied on humans.

## Acknowledgments

We are grateful to Peter W. Dicke and Friedemann Bunjes for technical support. This work was supported by the German Research Foundation (Deutsche Forschungsgemeinschaft) Grant FOR 1847-A3 (TH 425/13-1), received by P.T. within the framework of the Research Unit “Primate Systems Neuroscience,” and the Werner Reichardt Centre for Integrative Neuroscience, an excellence cluster grant received by the University of Tübingen (DFG EXC 307), coordinated by P.T.

